# Women born to older mothers have reduced fertility: Evidence from a natural fertility population

**DOI:** 10.1101/2020.09.18.303602

**Authors:** Niels van den Berg, Ingrid van Dijk, Rick Mourits

**Affiliations:** Leiden Universitair Medisch Centrum (LUMC), The Netherlands; Lund University, Sweden; Utrecht University, The Netherlands

**Keywords:** Fertility, mortality, reproductive ageing, mutation load, maternal age, family demography, fertility outcomes, neonatal mortality, age at last birth, reproductive senescence

## Abstract

Are daughters of older mothers less fertile? The human mutation rate is high and increases with chronological age. As female oocytes age, they become less functional, reducing female chances at successful reproduction. Increased oocyte mutation loads at advanced age may be passed on to offspring, decreasing fertility among daughters born to older mothers. In this paper we study the effects of maternal ageing on her daughter’s fertility, including total number of children, age at last birth, and neonatal mortality among her children. We study fertility histories of two generations of women from disjoint families from a pre-transitional historical population in the Dutch province of Zeeland. Using mixed effect Poisson models to take within family (sibling) relations into account, we show that fertility is reduced among married daughters who were born at advanced maternal age, with fewer children ever born and earlier ages at last birth. We do not find consistent evidence for effects on neonatal mortality. These results may indicate that women born to older mothers are negatively affected by their mothers’ increased oocyte mutation load.

## Introduction

Does a mother’s advanced age disadvantage her daughters’ reproduction? In the past decades, the age at first childbirth has increased. Understanding the consequences of late reproduction in humans is important because of the increasing postponement of childbearing (Sobotka 2004). There are strong indications that children of older mothers do not fare as well as offspring of younger mothers. A wide range of animal studies shows that higher parental age at conception is related to increased obesity, worse health, and fewer offspring (Heidinger et al., 2016; Hernández et al., 2020; Schroeder et al., 2015). Similarly, for in humans, there are also indications that women born to older mothers are less healthy, more often obese, have higher mortality, have fewer offspring, and have a larger chance of staying childless than their peers (for an overview of the literature see Barclay & Myrskylä, 2016).

However, studying the extent to which these effects are related to maternal ageing in data on contemporary human populations is complicated. Children born to older mothers are likely to benefit from secular improvements in mortality rates related to developing medical and public health conditions and educational expansion (Barclay & Myrskylä, 2016). In addition, in populations after the demographic transition later-born children are more likely to be born to higher-educated mothers. Furthermore, the total number of children today is strongly affected by the use of contraceptives, and late maternal reproduction is related to assisted reproductive technologies (ART). As such, for modern populations, establishing the relationship between being born to an older mother and own fertility is biased by many unobserved characteristics. Studying the consequences of late reproduction in humans requires multigenerational data from natural fertility populations, before socioeconomic differences in fertility or secular gains in lifespan, public health, and education properly set in.

We contribute to the literature on the influence of maternal age on her offspring by examining daughter’s reproduction patterns using a historical demographic database. We comprehensively examine the fertility of women born to older mothers as well as their children’s survival chances in a natural fertility population. Studying reproductive outcomes for older women’s daughters is relevant in its own right, as it can shed light on mechanisms of reproductive ageing. Additionally, studying reproductive health provides indications about women’s health for a period of life for which mortality is relatively low and other health outcomes can only rarely be studied (Van Dijk, Nilsson & Quaranta 2020). Finally, studying the trade-off between female reproduction at advanced ages and their offspring’s fertility helps us understand the association between ageing and biological fitness, which are concepts in evolutionary biology (Hernández et al., 2020).

We use a large historical database on the Dutch province of Zeeland (Mandemakers & Laan, 2017) to study natural fertility birth cohorts, allowing us to study the total number of children instead of age at menopause, which is not necessarily affected by mutation load and oocyte quality. Life courses of women and their mothers are observed before the demographic transition, implying that the fertility patterns observed are not the product of conscious fertility control and that fertility and mortality patterns remain constant over time. We build upon the work of Gillespie, Russell and Lummaa (2013), who showed that older mothers negatively affect offspring survival and fertility in four historical Finnish parishes. Although these first results are in line with empirical observations from animal studies, the statistical estimates are not robust due to the small sample size. Therefore, we use historical data on an entire province to assess whether and to what extent time to conception, total number of children, and neonatal mortality are related to the age of the grandmother at the time of birth of the mother.

### Data construction and analytical strategy

The LINKS data contains information on individuals who were born, married or died in the Dutch province of Zeeland between 1812 and 1957. Civil certificates have been linked together into life courses and family trees, containing up to nine generations of relatives. A more extensive description of the data and the data quality may be found elsewhere (Van den Berg et al., 2020; Mourits, Van Dijk & Mandemakers, 2020). From these data, we select three-generation families (see Figure 1). The first generation, the mothers (F0), consists of women who married between 1812-1850. Mothers and their first spouse survived to her 50^th^ birthday or older, and they had at least one child. Research persons (RPs, F1) are born between 1812 and 1874 and had their last children (F2) before 1914, with few births after 1900, so that results are not affected by the general decline of the number of births in the province of Zeeland (the demographic transition). In total we identified 4,428 mutually exclusive three generation families covering 7,332 F0 parents, 9,732 F1 RPs, and 73,075 F2 children.

**FIGURE 1.**
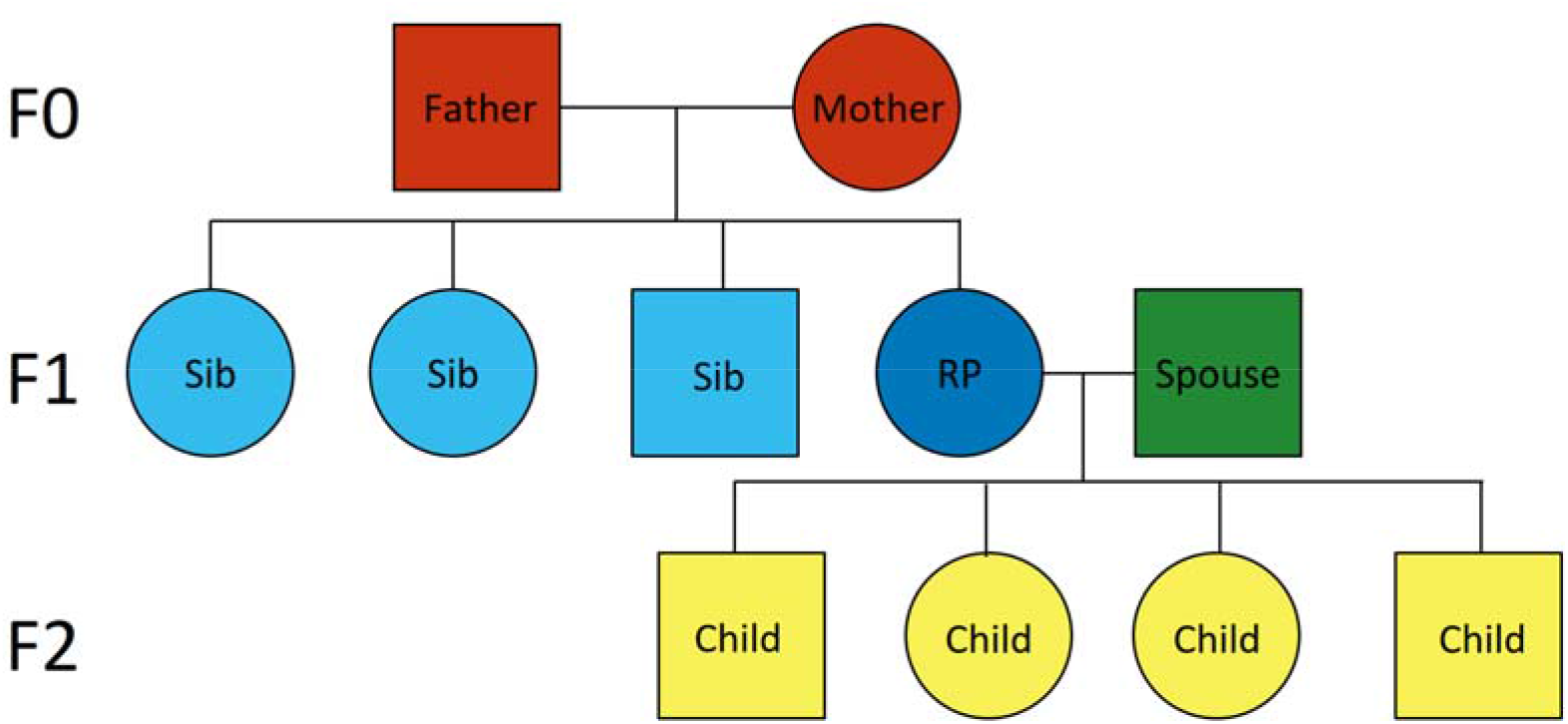
Example family tree of the selected research persons (RPs)

**TABLE 1.** Descriptive statistics

|  | F0 - Mothers | F1 - Research persons | F2 - Children |
| --- | --- | --- | --- |
| Number (N) | 7,332 | 9,732 | 7,3075 |
| Deceased (%) | 6,362 (87%) | 9,732 (100%) | 5,3301 (73%) |
| Missing age (%) | 1 (<0%) | 0 (0%) | 25 (0%) |
| Censored (%) | 969 (13%) | 0 (0%) | 19,749 (27%) |
| Mean age at last observation | 60.03 (18.23) | 73.9 (10.43) | 30.15 (31.3) |
| Mean age at death | 63.41 (17.07) | 73.9 (10.43) | 34.33 (34.48) |
| Females (%) | 7,332 (100%) | 9,732 (100%) | 35,441 (48%) |
| Birth cohorts | 1768-1833 | 1812-1874 | 1832-1914 |
| Marriage cohorts | 1812-1850 | 1831-1905 | 1852-1937 |
| Mean number of children | 7.66 | 7.51 | 4.89 |

## Results

Results are presented in Table 2.

**TABLE 2.** Results of models of fertility careers of women (F1) born to older mothers (F0)

|  | **Model 1:**<br>**Age mother at birth RP** | **Model 2:**<br>**Marriage age** | **Model 3:**<br>**Birth year mother** |
| --- | --- | --- | --- |
| **Number of children** |  |  |  |
| N (mean) | 9732 (7.508) |  |  |
| IRR (95% CI) | 0.997 (0.996-0.999) | 0.941 (0.939-0.943) | 0.997 (0.997-0.998) |
| p-Value | $2.38*10^{-3}$ | $<2.00*10^{-16}$ | $1.21*10^{-7}$ |
| **Neonatal mortality** |  |  |  |
| N (mean) | 9732 (0.715) |  |  |
| IRR (95% CI) | 0.995 (0.990-1.00) | 0.980 (0.973-0.987) | 0.993 (0.990-0.996) |
| p-Value | $5.41*10^{-2}$ | $4.11*10^{-8}$ | $4.16*10^{-6}$ |
| **Age last birth** |  |  |  |
| N (mean) | 9732 (39.616) |  |  |
| IRR (95% CI) | 0.975 (0.960-0.989) | 1.193 (1.168-1.220) | 0.970 (0.962-0.979) |
| p-Value | $1.01*10^{-3}$ | $<2.00*10^{-16}$ | $1.46*10^{-11}$ |

Table 2 presents the effect of age of the mother at birth RP on her number of children, neonatal mortality, and age at last birth, respectively. Model 1 is the uncontrolled effect, in model 2 marriage age is controlled for, and in model 3 birth year of the mother is controlled for.

We estimate a Poisson distributed generalized linear mixed model to study the relation between mother age at birth and the count distribution of 1) number of children, 2) neonatal mortality, and 3) age at last birth. We model a random effect to adjust for within-family relations of the F1 PRs. Specifically, we allow a random intercept per family so that we study within family (within RP sibships, clustered by their mother) effects.

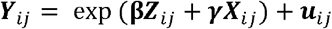

Where ***Y**_ij_* is vector of responses for the j-th F1 RP of F0 mother i and corresponds to either the number of children, neonatal mortality, or age at last birth. **β** is a vector of regression coefficients for the main effects of interest (***Z***) which corresponds to the age of the mother at the time of birth of the RP, ***γ*** is a vector of regression coefficients for the effects of covariates and possible confounders (***X***), which are marriage age, and birth year of the mother. ***u*** refers to an unobserved random effect shared by RPs of a given mother. In contrast to work by Gillespie, Russell, and Lummaa (2013), the models are adjusted for marriage age of the mother rather than her total number of children, as her marriage age is a more precise estimate for the period in which she could have had children and may be affected by parental socioeconomic resources in this historical population. Furthermore, results are controlled for possible trends over time by including the birth year of the RP’s mother rather than the birth year of her children, so that trends over time and the ageing of the mother are included as two separate effects.

The results provide evidence that daughters of older mothers have a decreased fertility. In Table 2 we show that the incidence rate ratio (IRR) for number of children is 0.997. This indicates that for every unit increase in the age of RP’s mother at birth of the RP the incidence rate of number of children decreases by a factor of 0.997 or, in other words, decreases with 0.3%. The effect is thus robust but small. Furthermore, we find that the age at last birth is lower for women with older mothers. For each yearly increase in the age at birth of the mother, her daughter’s age at last birth decreased with 9 days. The effect is significant. Finally, the neonatal mortality is lower among women with older mothers, but the effect is not very precisely estimated and marginally insignificant (p=0.054). Additional analyses (tables can be provided upon request) show that the effect on neonatal mortality is limited to the earlier cohorts and disappeared when child mortality started declining and less than 30% of the population died in first year of life. Neonatal mortality consequences of motherhood at advanced ages are likely to be even lower in modern low-mortality contexts, but there is little reason to assume that fertility consequences among women with older mothers have dissipated over time.

## Discussion and conclusion

We presented evidence that late maternal reproduction is associated with multiple negative health outcomes in her daughters in a natural fertility population in Zeeland, The Netherlands, 1812-1874. As the age of their mother increased, women’s number of children decreased and their age at last birth declined. These outcomes are in line with earlier studies which showed that late reproduction is associated with decreased reproduction in various animals and an earlier age at menopause in humans. It has been hypothesized that one of the main drivers behind this effect is aging of the germ cells (reproductive ageing) (Cawthon et al. 2020). Over the life time mutations, both in somatic and germ cells, accumulate, albeit at a much slower rate in germ cells. These mutations are associated to a decreased fertility and health (Holmes et al., 1992). Older mothers’ higher germ cell mutation rate is possibly related to a higher germline mutation rate in their children (Cawthon et al., 2020; Schroeder et al, 2015). Higher germline mutation rates related to a reduced reproduction for later-born children are consistent with our findings. Earlier research may also provide an alternative explanation for our findings as it has shown that with age the length of telomeres decreases. Telomeres are the protective ends of chromosomes and shorten with every cell division until the telomere is critically shortened and the cell dies. Telomere length is therefore strongly associated with chronological age, including age related diseases. There is some evidence from animal studies that telomere length is heritable. As a consequence, daughters of older mothers may have shorter telomeres, which may be related to accelerated ageing and reproductive ageing, possibly resulting in reduced fertility (Schroeder et al., 2015).

The association between maternal age at birth and offspring fertility is probably not only caused by effects of reproductive ageing, but also by effects of earlier reproduction, the composition of the family network, and due to their relative robustness. In Zeeland, older mothers not only reproduced until advanced ages, but also gave birth to more children during their lifetime. Due to nutritional deficiencies and short birth intervals, women had few resources and little time to physically recover from previous pregnancies, a phenomenon known as maternal depletion. The physical depletion of mothers is related to decreased birth weight among their children, which made them more vulnerable in the first years of life and is related to a range of negative life course outcomes.

Second, women who have older mothers have, on average, different family networks than their earlier-born siblings. They are more likely to lose their parents at a young age, especially in historical populations with shorter adult life expectancy, which increases their mortality and may hasten their transition into marriage (Rosenbaum-Feldbrügge, 2020; Störmer & Lummaa, 2014; Voland & Willführ, 2017). When daughters marry and have children themselves, later-born children in the sibship have older parents who are less likely to be alive and help raise their grandchildren. A body of earlier research has suggested that especially maternal grandmothers contribute positively to their daughter’s reproduction and infant and child survival (Hawkes, 2013; Sear & Mace 2008). As such, women with older mothers may be relatively disadvantaged when their children are young and neonatal mortality among their children could be relatively higher, due to differences in their family networks.

A study on a historical population from Utah showed that mortality rates are decreased for children born to mothers with the capacity to reproduce at advanced ages, conditional on her survival to age 50 (Hin et al. 2016). As such, mothers who are able to have children at an advanced age may be relatively robust and their survival advantage, on average, may be shared with their children. At the same time, their relative robustness may not be shared with all their children equally. As we have illustrated in the current work, late-born children may be subject to patterns of accelerated (reproductive) ageing and decreased fertility. Whether ageing and survival at later ages are affected as well remains an open question.

In sum, mother age at birth has a small, but wide-ranging effect on her offspring’s life course. Daughters of older mothers had fewer children of their own, and had their last child at an earlier age. These insights from 19^th^ century Zeeland are still very much relevant today. Reproduction is more limited today and subject to deliberate birth control, but since the 1970s the average age at last birth has increased and late-age reproduction has become more common than in the mid-century. However, late reproduction is not a historical exception, as women had children throughout their fertile window in recent history. At the same time, we have shown evidence that having children at advanced ages may come at a price. The effects of maternal age on offspring fertility may still have a small but significant impact on fertility of women today.

In future work, further disentanglement of the impact of maternal age on children’s reproduction should take fathers into account next to mothers. Next to women, men’s reproductive career could be affected by parental mutation accumulation indicated by mother’s advanced age at their birth (Arslan et al., 2017; Hayward, Lummaa & Bazykin, 2015). Furthermore, marital success may play a key role in linking together mother’s age and her children’s own fertility, both from a social and a biological perspective. Children of older mothers, and to some extent also fathers. are more prone to have health syndromes, such as Down’s Syndrome, and other physical deficiencies, which could lead to lower marital and reproductive success. Children of older parents may be disadvantaged on the marriage market also due to caretaking burdens for their parents (Hayward, Lumma & Bazykin, 2015). Evidence presented here concerns married individuals, and more wide-ranging effects of advanced maternal age on reproduction may be identified taking into account a broader selection of her children, but is beyond the scope of this paper. To conclude, the ramifications of late reproduction on offspring health are much broader than the presented evidence that reproduction and reproductive ageing are affected by maternal age at birth and socio-biological frameworks are necessary to understand why.

## Acknowledgements

We are greatly indebted to Kees Mandemakers for his indispensable labor on the HSN and LINKS projects. These projects made it possible to study life courses, familial ties, intergenerational effects, and demographic patterns for the 19^th^ and 20^th^ century Netherlands. First, in a sample of the Dutch historical population, then as the entire historical population of the province of Zeeland, and in the near future of The Netherlands. These projects have radically improved historiography related to the social, economic and demographic developments in the history of The Netherlands. The sources are an indispensable source for future historical, demographic, medical, and social research. Work such as presented here would not be possible without his efforts to advance our knowledge of the Dutch population history. We thank Kees for his efforts throughout his career to advance the project.

## Notes

### Competing Interest Statement

The authors have declared no competing interest.

## References

Barclay, K., & Myrskylä, M. (2016). Advanced Maternal Age and Offspring Outcomes: Reproductive Aging and Counterbalancing Period Trends. Population and Development Review, 42(1), 69–94. JSTOR.

Berg, N. van den, Dijk, I. K. van, Mourits, R. J., Slagboom, P. E., Janssens, A. A. P. O., & Mandemakers, K. (2020). Families in comparison: An individual-level comparison of life-course and family reconstructions between population and vital event registers. Population Studies, 25(3), 484–526. https://doi.org/10.1080/00324728.2020.1718186

Cawthon, R. M., Meeks, H. D., Sasani, T. A., Smith, K. R., Kerber, R. A., O’Brien, E., Baird, L., Dixon, M. M., Peiffer, A. P., Leppert, M. F., Quinlan, A. R., & Jorde, L. B. (2020). Germline mutation rates in young adults predict longevity and reproductive lifespan. Scientific Reports, 10(1), 10001.

Dijk, I.K. van, Nilsson & Quaranta. Unpublished paper. Disease exposure in early life affects female reproduction: Evidence from Southern Sweden 1896–2000

Gillespie, D. O. S., Russell, A. F., & Lummaa, V. (2013). The Effect of Maternal Age and Reproductive History on Offspring Survival and Lifetime Reproduction in Preindustrial Humans. Evolution, 67(7), 1964–1974.

Hawkes, K., & Coxworth, J. E. (2013). Grandmothers and the evolution of human longevity: a review of findings and future directions. Evolutionary Anthropology: Issues, News, and Reviews, 22(6), 294–302.

Hernández, C. M., Daalen, S. F. van, Caswell, H., Neubert, M. G., & Gribble, K. E. (2020). A demographic and evolutionary analysis of maternal effect senescence. Proceedings of the National Academy of Sciences, 117(28), 16431–16437.

Holmes, G. E., Bernstein, C., & Bernstein, H. (1992). Oxidative and other DNA damages as the basis of aging: A review. Mutation Research/DNAging, 275(3), 305–315.

Schroeder, J., Nakagawa, S., Rees, M., Mannarelli, M.-E., & Burke, T. (2015). Reduced fitness in progeny from old parents in a natural population. Proceedings of the National Academy of Sciences, 112(13), 4021–4025.

Sear, R., & Mace, R. (2008). Who keeps children alive? A review of the effects of kin on child survival. Evolution and human behavior, 29(1), 1–18.

Sobotka, T. 2004. Postponement of childbearing and low fertility in Europe. University of Groningen, The Netherlands.

Arslan, R.C., Willführ, K.P., Frans, E.M., Verweij, K.J.H., Bürkner, P-C., Myrskylä, M., Voland, E., Almqvist, C., Zietsch, B.P. & Penke, L. (2017). Older fathers’ children have lower evolutionary fitness across four centuries and in four populations. Proceedings of the Natural Society B: Biological Sciences, 284(1862), 20171562.

Hayward, A.D., Lummaa, V. & Bazykin, G.A. (2015). Fitness consequences of advanced ancestral age over three generations in humans. PLoS ONE, 10(6), e0128197.

Heidinger, B.J., Herborn, K.A., Granroth-Wilding, H.M.V., Boner, W., Burthe, S., Newell, M., Wanless, S., Daunt, F., Monaghan, P. (2015). Parental age influences offspring telomere loss. Functional Ecology, 30, 1531–1538.

Hin, S., Ogórek, B., & Hedefalk, F. (2016). An old mom keeps you young: Mother’s age at last birth and offspring longevity in nineteenth-century Utah. Biodemography and Social Biology, 62(2), 164–181.

Mandemakers, K., & Laan, F. (2017). LINKS dataset Genes Germs and Resources, WieWasWie Zeeland, Civil Certificates, version 2017.01 [Data file and code book]. Amsterdam: IISH.

Mourits, R.J., Van Dijk, I.K. & Mandemakers, K. (2020). From matched certificates to related persons: Building a dataset from LINKS-Zeeland, Historical Life Course Studies, forthcoming.

Rosenbaum-Feldbrügge, M. (2020). Dealing with demographic stress in childhood: Parental death and the transition to adulthood in the Netherlands, 1850–1952. Enschede: Ipskamp.

Störmer, C., & Lummaa, V. (2014). Increased mortality exposure within the family rather than individual mortality experiences triggers faster life-history strategies in historic human populations. PloS One, 9(1), 1–9.

Voland, E., & Willführ, K. P. (2017). Why does paternal death accelerate the transition to first marriage in the C18-C19 Krummhörn population? Evolution and Human Behavior, 38(1), 125–135.

